# Dissecting Rate-Limiting Processes in Biomolecular Condensate Exchange Dynamics

**DOI:** 10.1101/2025.05.16.654578

**Authors:** Ross Kliegman, Eli Kengmana, Rebecca Schulman, Yaojun Zhang

**Author notes:** (R.K.). Contact authors (Y.Z.).

## Abstract

An increasing number of biomolecules have been shown to phase-separate into biomolecular condensates — membraneless subcellular compartments capable of regulating distinct biochemical processes within living cells. The speed with which they exchange components with the cellular environment can influence how fast biochemical reactions occur inside condensates and how fast condensates respond to environmental changes, thereby directly impacting condensate function. While Fluorescence Recovery After Photobleaching (FRAP) experiments are routinely performed to measure this exchange timescale, it remains a challenge to distinguish the various physical processes limiting fluorescence recovery and identify each associated timescale. Here, we present a reaction-diffusion model for condensate exchange dynamics and show that such exchange can differ significantly from that of conventional liquid droplets due to the presence of a percolated molecular network, which gives rise to different mobility species in the dense phase. In this model, exchange can be limited by diffusion of either the high- or low-mobility species in the dense phase, diffusion in the dilute phase, or the attachment/detachment of molecules to/from the network at the surface or throughout the bulk of the condensate. Through a combination of analytic derivations and numerical simulations, we quantify the contributions of these distinct physical processes to the overall exchange timescale and predict an experimentally testable scaling relationship between the exchange timescale and condensate size. We discover that the exchange dynamics can be accelerated via a pore-mediated pathway in which molecules pass through the pores of the meshwork and attach/detach directly in the condensate interior. Notably, this pathway leads to a new regime in which the exchange timescale becomes independent of condensate size, which we validate through FRAP experiments on a biosynthetic DNA nanostar system. Our work offers insight into the predominant physical mechanisms driving condensate material exchange, with implications for natural and engineered systems.

## INTRODUCTION

Living cells organize their contents into distinct functional compartments. Beyond traditional membranebound organelles, subcellular structures can take the form of dynamic, liquid-like networks of molecules called “biomolecular condensates” [1, 2]. These condensates are dense assemblies of distinct proteins and nucleic acids that are driven by multivalent interactions to segregate out of the intracellular milieu. They enable functions vital for life, including gene regulation [3–5], signal transduction [6–8], and stress response [9–11], and when misregulated, they have been implicated in various diseases, most notably neurodegeneration [12–14] and cancer [15–18]. Understanding how condensates form and evolve over time in cells can therefore deepen our physical understanding of emergent self-organization in biological systems and potentially inform human health.

The liquid-like nature of condensates is often critical for their biological function, as it enables exchange of material with the surrounding dilute phase. For instance, metabolic condensates, such as purinosomes [19, 20], are enriched in enzymes, substrates, and other biomolecules involved in specific metabolic pathways [21]. Regulating metabolic activity in condensates requires not only that reactants can partition into them, but also that products can later escape. However, with viscosities orders of magnitude larger than conventional oil droplets [22], condensates are thought to experience slow internal diffusion, limiting the exchange dynamics. More broadly, the speed of material exchange can influence condensate responses to environmental changes, as well as the number, size, and spatial distribution of condensates via Ostwald ripening [23, 24]. Collectively, these effects can impact condensate function, motivating a need for tools to accurately measure and interpret the exchange dynamics.

The timescales of molecular exchange are commonly measured with an experimental technique known as Fluorescence Recovery After Photobleaching (FRAP) [25–28]. In a typical FRAP experiment, fluorescently labeled molecules are photobleached within a region of interest (ROI) upon irradiation with a high-intensity laser. The fluorescence intensity in the ROI then recovers over time due to molecular exchange with the surroundings until constant intensity is eventually restored. Photobleaching can be performed on a subregion within a droplet, known as partial FRAP, or on an entire droplet, known as full FRAP. Whereas partial FRAP experiments predominantly measure internal mixing within the dense phase, exchange between dense and dilute phases can be probed using full FRAP. Exchange dynamics have been studied in a range of experimental condensate systems [6, 10, 29– 32], and complementary theories have been developed to extract meaningful physical quantities from measured fluorescence recovery curves [33–36]. While it is typically assumed that the exchange dynamics are limited by molecular diffusion, recent studies suggest that condensate material exchange can also be limited by other physical processes owing to the complexity of molecular interactions [37–41], e.g., interface resistance [40, 41].

The exchange dynamics of condensates are ultimately determined by the constituent biomolecules and their microscopic structures and interactions. These interactions often include a complex combination of electrostatic, *ππ*, cation-*π*, hydrophobic, hydrogen-bonding, and complementary domain-domain interactions [42, 43]. It has been proposed that a range of molecules, from intrinsically disordered proteins with low-complexity sequences to associative proteins with multiple interaction domains, generally conform to a “sticker-spacer” architecture [44–46], in which “stickers” are residues, sequence motifs, or larger folded domains that form reversible physical cross-links driving phase separation, and “spacers” exclude volume and connect the stickers to form polymers.

In the sticker-spacer framework, it follows that phase-separating molecules often form dynamically restructuring networks that go beyond traditional liquid-liquid phase separation (Fig. 1a), sometimes referred to as “phase separation coupled to percolation” [47, 48]. In the modified physical picture (Fig. 1b), attachment and detachment of molecules to and from the percolated network naturally gives rise to different mobility populations within the condensate for the same type of molecule. The low-mobility population (referred to as “species 1”) represents molecules bound to the network, and the high-mobility population (referred to as “species 2”) represents freely diffusing molecules detached from the network.

**FIG. 1.**
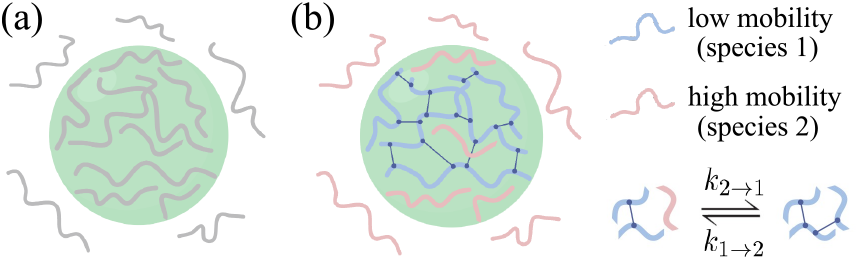
Schematics of a condensate in (a) the conventional model, which assumes uniform molecular mobility inside the condensate (depicted in grey), and (b) our proposed model, in which transient attachment to/detachment from the molecular network can give rise to multiple mobilities for the same molecule inside the condensate. Connected blue molecules are bound to the network, whereas individual pink molecules are freely diffusing. By attaching and detaching, the two mobility species can convert between one another with rates *k*_2→1_ and *k*_1→2_, respectively.

Indeed, multiple mobility populations have been reported in the dense phase of an *in vitro* reconstituted postsynaptic density system [49] as well as in single-component A1-LCD condensates [50]. Specifically, in the postsynaptic density system, adaptive single-molecule tracking has revealed that molecules within condensates stochastically switch between confined and freely diffusing states, consistent with motion on and off a percolated molecular network [49]. In A1-LCD condensates, single-fluorogen imaging has revealed that slow fluorogens localize to nanoscale hydrophobic hubs, whereas faster fluorogens explore more weakly interacting regions of the condensate, supporting the existence of a spatially heterogeneous landscape for condensate dynamics [50]. Together, these studies demonstrate that even nominally single-component condensates can harbor rich internal dynamical heterogeneity, yet a theory to interpret such dynamic complexity has been lacking.

Here, we develop a minimal reaction-diffusion model that incorporates two mobility species in the dense phase. We show that exchange can be accelerated through a pore-mediated pathway, where molecules pass through meshwork pores and attach/detach directly within the condensate interior. Using theory and simulations, we identify distinct rate-limiting processes with characteristic scaling relationships, and validate these predictions with FRAP experiments on DNA nanostar condensates.

## RESULTS

### A. reaction-diffusion model for condensate exchange dynamics

To explore how the presence of two mobility species impacts the condensate exchange timescale, we first develop a reaction-diffusion model for a phase-separated system at equilibrium. Assuming spherical symmetry, we describe the recovery dynamics of a fully bleached condensate (equivalent to the exchange dynamics) by the following coupled reaction-diffusion equations:

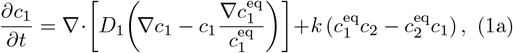

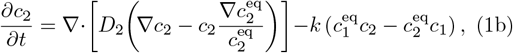

where *c*_1_(*r, t*) and *c*_2_(*r, t*) are the bleached concentrations of species 1 and 2, respectively, *D*_1_(*r*) and *D*_2_(*r*) are their position-dependent diffusion coefficients, 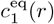 and 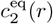 are their equilibrium concentration profiles, and *k* is a parameter that encodes how fast molecules convert between species. The coordinate *r* is the distance from the center of the spherical condensate, and *t* denotes the time.

In Eqs. (1a) and (1b), the first terms on the right-hand side represent conventional Fickian diffusion in a concentration gradient, and the second terms represent excess chemical potentials that drive molecules towards nonuniform equilibrium concentration profiles [34]. The third and fourth terms account for mobility switching due to binding/unbinding with the network. Molecules can attach to the percolated network and lower their mobility with a rate *k*_2→1_(*r*), and detach from the network and regain higher mobility with a rate *k*_1→2_(*r*). Detailed balance requires that the fluxes of association (*c*_2_*k*_2→1_) and dissociation (*c*_1_*k*_1→2_) are equal at equilibrium, which allows us to characterize these rates in terms of a single parameter,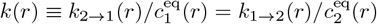. For simplicity, we assume *k* to be a constant, independent of the distance from the center of the condensate. The association and dissociation fluxes are then 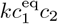 and 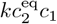, respectively. We note that only the free form of scaffold molecules is present in the dilute phase with no percolated network, i.e., 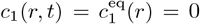 (later referred to as *c*_1,dil_ = 0). Consequently, there is no association or dissociation flux outside the condensate, and the dynamics of free scaffold molecules are described by a single Eq. (1b) without the third and fourth terms on the right.

Analogous to a full FRAP experiment, we set the initial condition to be

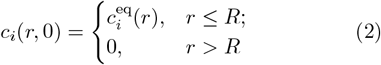

for a fully bleached droplet, where *i* = 1, 2, and *R* is the droplet radius. We impose no-flux boundary conditions to conserve total particle number in the system:

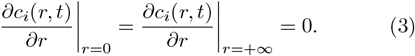

Upon solving for *c*_1_(*r, t*) and *c*_2_(*r, t*), we can obtain a normalized brightness curve *I*(*t*) for the fraction of molecules inside the droplet that are unbleached at a time *t*:

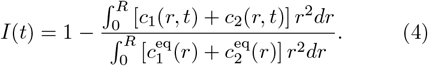

Finally, the characteristic timescale *τ* of the exchange dynamics is identified by fitting *I*(*t*) to an exponential function of the form 1 − *e*^−*t/τ*^ . We note that the system reaches equilibrium when 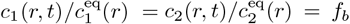, where the constant *f*_*b*_ is the fraction of total molecules that are bleached. In most cases, *f*_*b*_ ≪ 1, so that *I*(*t* → ∞) = 1.

### Quantifying the timescales of rate-limiting processes

#### Analytical derivations

A proxy for condensate material exchange, fluorescence recovery in a bleached droplet is a multi-step process involving dilute-phase diffusion, network attachment/detachment, and dense-phase diffusion. We outline the rate-limiting steps of FRAP recovery in Fig. 2. First, unbleached molecules must diffuse through the dilute phase to reach the droplet surface. In the limit of low dilute-phase concentration, we derive the dilute-phase diffusion-limited timescale *τ*_dil_ shown in Eq. (5a). Next, in the absence of species 2 in the dense phase (i.e., *c*_2,den_ = 0), unbleached molecules will enter the droplet by attaching to the network at the droplet interface and subsequently diffuse into the bulk dense phase as species 1. In this case, if interfacial attachment/detachment is rate-limiting, we derive the interface-limited timescale *τ*_int_ shown in Eq. (5b), whereas if dense-phase diffusion of species 1 is rate-limiting, we derive the timescale *τ*_1,den_ shown in Eq. (5c):

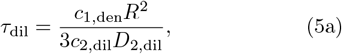

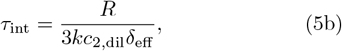

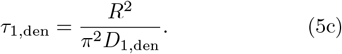

*δ*_eff_ is the effective width of the droplet interface, and *c*_*i*,dil_ and *D*_*i*,dil_ (*c*_*i*,den_ and *D*_*i*,den_) are the equilibrium concentration and diffusion coefficient of species *i* in the dilute (dense) phase, respectively, with *i* = 1, 2. These timescales largely follow our previous results in [40]. Detailed derivations of each timescale are provided in the Supplemental Material [51].

**FIG. 2.**
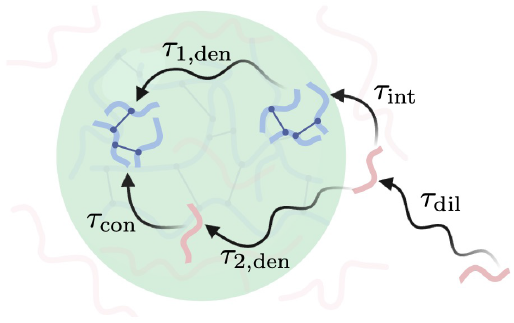
Schematic of rate-limiting processes in the exchange dynamics of biomolecular condensates. In order for fluorescence to recover in a bleached droplet, unbleached molecules have to diffuse in the dilute phase with a timescale *τ*_dil_ until they encounter the droplet, attach to the network at the interface with a timescale *τ*_int_ or within the droplet bulk with a timescale *τ*_con_, and diffuse within the network as species 1 (blue) with a timescale *τ*_1,den_ or through the network mesh inside the droplet as species 2 (pink) with a timescale *τ*_2,den_. In the slow attachment limit, the faster of the two attachment mechanisms (*τ*_int_ vs. *τ*_con_) controls the exchange speed, whereas in the slow dense-phase diffusion limit, the faster of the two dense-phase diffusive mechanisms (*τ*_1,den_ vs. *τ*_2,den_) controls the exchange speed.

In the presence of species 2 in the dense phase (i.e., *c*_2,den_ > 0), a pore-mediated mode of recovery emerges as unbleached molecules can now alternatively enter the droplet by passing through the pores of the network and diffusing into the dense phase as species 2, which then attach to and detach from the network throughout the bulk of the droplet (Fig. 2). If the conversion between species 1 and 2 is fast relative to the diffusion timescales (i.e., *k* is sufficiently large), the bleached concentrations *c*_1_ and *c*_2_ can be treated as having reached local equilibrium: 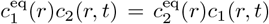 Eqs (1a) and (1b) then reduce to a single “effective diffusion” equation:

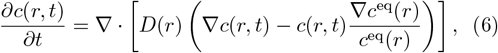

where *c*(*r, t*) = *c*_1_(*r, t*) + *c*_2_(*r, t*) is the total bleached concentration profile, 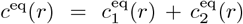 is the total equilibrium concentration profile, and [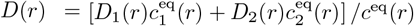] is the effective diffusivity profile. In this fast-conversion regime, if the dilute-phase diffusion is rate-limiting, the recovery timescale reverts to *τ*_dil_ in Eq. (5a) with *c*_1,den_ replaced by *c*_1,den_ + *c*_2,den_. However, if the dense-phase diffusion is rate-limiting, we derive a new recovery timescale:

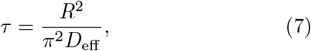

where *D*_eff_ is the effective dense-phase diffusion coefficient [52, 53]:

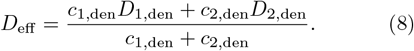

Since molecules can either attach and diffuse as species 1 or detach and diffuse as species 2, *D*_eff_ is a weighted sum of the two diffusion processes, with the weights given by their respective occupancies. By definition, species 2 molecules diffuse more rapidly than species 1 (i.e., *D*_2,den_ ≫ *D*_1,den_). Thus, the presence of species 2 molecules in the dense phase can speed up the exchange dynamics (i.e., *D*_eff_ > *D*_1,den_). However, because molecules may spend much less time as species 2 than as species 1, the degree of acceleration depends on the specific parameter values. When diffusion as species 1 dominates (i.e., *D*_1,den_ ≫ *D*_2,den_*c*_2,den_/*c*_1,den_), the recovery timescale is comparable to *τ*_1,den_ in Eq. (5c). Conversely, when species 2 diffusion dominates, the acceleration is significant and the timescale becomes:

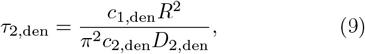

where we have assumed *c*_1,den_ ≫ *c*_2,den_ (justified below). Detailed derivations are provided in the Supplemental Material [51].

In the opposite limit, if the diffusive processes are much faster than the conversion, then the bleached molecules of species 2 inside the droplet will be quickly replaced by unbleached molecules diffusing in from the dilute phase. This leads to a fast and partial recovery, with the recovery amplitude determined by the fraction of species 2 molecules in the dense phase [*c*_2,den_/(*c*_1,den_ + *c*_2,den_)]. In contrast, the bleached molecules of species 1 can only be replaced via a slow attachment/detachment process, where they exchange with unbleached molecules of species 2 at the droplet interface and throughout the droplet bulk. This leads to a slow mode of recovery following the initial fast mode until the droplet is fully recovered. We show an example of such two-step recovery at the end of the following subsection (*Numerical simulations*). Here, we argue that in many cases, species 1 is expected to be much more abundant than species 2 in the dense phase (i.e., *c*_1,den_ ≫ *c*_2,den_) because species 1 is bound to the network and therefore energetically favored over species 2. In this situation, the contribution of the first fast mode is negligible so that the recovery curve mainly reflects the second slow process. We will thus focus on the recovery timescale of this second conversion-limited mode in the following.

Since conversion is slower than diffusion, the bleached molecules of each species can be considered equilibrated with respect to their equilibrium concentration profiles:

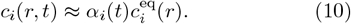

The time-dependent component of *c*_*i*_(*r, t*) is given by *α*_*i*_(*t*), which asymptotically approaches the overall bleached fraction, *f*_*b*_, for *i* = 1, 2. To derive the timescale of recovery, we first apply the divergence theorem to Eqs. (1a) and (1b), yielding:

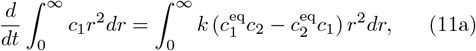

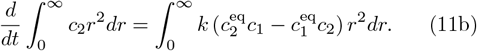

We then substitute the approximate forms of *c*_1_(*r, t*) and *c*_2_(*r, t*) [Eq. (10)] into Eq. (11), leading to a relaxation time for *α*_1_(*t*) and *α*_2_(*t*):

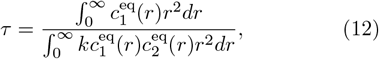

which is the conversion-limited recovery timescale.

There are two contributions to the integral in the denominator of Eq. (12): the integration over the interface region and the integration throughout the bulk of the condensate. By separating them into two terms, we derive in the Supplemental Material [51] an explicit form of the conversion-limited timescale:

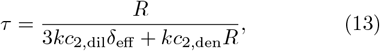

which combines the effects of network attachment both at the interface and in the bulk of the droplet. We note that we have assumed a constant conversion rate *k* at the interface and in the condensate bulk. If the conversion rates differed in these two regions, the region-specific rates *k*_int_ and *k*_con_ would replace *k* in the interface- and conversion-limited contributions to the denominator, respectively. Once again, the presence of species 2 molecules in the dense phase can speed up the exchange dynamics, with the degree of acceleration determined by the specific parameter values. When conversion at the interface dominates the recovery time (i.e., *c*_2,dil_*δ*_eff_ ≫ *c*_2,den_*R*), the recovery timescale is comparable to *τ*_int_ in Eq. (5b). Conversely, when conversion throughout the condensate bulk dominates, the acceleration becomes significant and we obtain the bulk conversionlimited timescale:

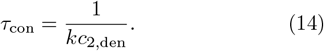

At this point, we have identified five rate-limiting processes for the exchange dynamics of droplets with two mobility species and derived analytical expressions for their corresponding timescales in Eqs. (5), (9), and (14). Each physical process has a distinct timescale that scales with droplet size differently. Intuitively, the diffusion-limited processes in both dense and dilute phases are associated with timescales that scale as *R*^2^/*D* [Eqs. (5a), (5c), and (9)]. The prefactors *c*_1,den_/*c*_2,dil_ and *c*_1,den_/*c*_2,den_ in Eqs. (5a) and (9) account for replacing bleached molecules of concentration *c*_1,den_ with unbleached molecules of concentrations *c*_2,dil_ and *c*_2,den_, respectively. The interfacial timescale [Eq. (5b)] accounts for exchange of a volume of molecules (∼ *R*^3^) over a surface (∼ *R*^2^) and is therefore linear in *R*. Lastly, the bulk conversion-limited timescale [Eq. (14)] is independent of *R*, which arises due to slow detachment of bleached molecules throughout the bulk of the dense phase. Once detached, these molecules can quickly escape the droplet, allowing unbleached molecules to attach to the network. The recovery timescale is therefore limited by the lifetime of a molecule in the network: 1/*k*_1→2_ = 1/(*kc*_2,den_).

Putting together the rate-limiting steps, we propose the following expression for the overall timescale of fluorescence recovery:

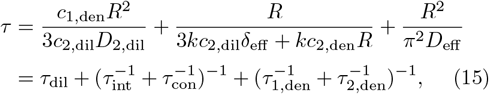

where following diffusion in the dilute phase, two additional processes, network attachment/detachment and dense-phase diffusion, are required for total fluorescence recovery. Again, we have invoked the assumption that *c*_1,den_ ≫ *c*_2,den_, which is motivated by the lower energy attained when molecules are attached to the droplet network. Both network attachment/detachment and dense-phase diffusion involve a combination of two competing processes in parallel: molecules can attach/detach at the interface as well as in the bulk of the condensate, and they can diffuse as species 1 or species 2, the slower of which is neglected in favor of the faster. It is worth noting that by setting *c*_2,den_ = 0 in Eq. (15), i.e., assuming a single mobility species inside the droplet, we recover results of our previous study [40]: *τ* = *τ*_dil_+*τ*_int_+*τ*_1,den_. In particular, *τ*_int_ arises due to the droplet’s “interface resistance”, which was previously modeled with a phenomenological parameter *κ* but now acquires a clear physical meaning: *τ*_int_ is governed by the molecular attachment/detachment at the droplet interface. For *c*_2,den_ > 0, the emergence of a pore-mediated mode of recovery in Fig. 2 can speed up the exchange dynamics and leads to two previously unrecognized terms in the recovery time, *τ*_con_ and *τ*_2,den_, resulting in a complex dependence of *τ* on the droplet radius *R*. We demonstrate this complex dependence via numerical simulations and FRAP experiments on DNA nanostar droplets below.

#### Numerical simulations

In the previous section, we derived the timescale of fluorescence recovery by analytically solving the reaction-diffusion system described by Eqs. (1-4) in various limits. Here, we numerically verify these timescales and visualize the different FRAP signatures in each rate-limiting case. Consistent with equilibrium solutions of the Cahn-Hilliard equation [54], we first specify the functional form of the equilibrium concentration profiles with a sharp but smooth transition at the droplet interface (*r* = *R*) over a finite width ∼ *l* (Fig. 3a):

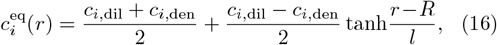

with *i* = 1, 2, and we assume a similar form for the diffusivity profiles (Fig. 3b):

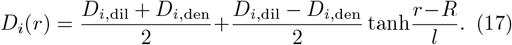

**FIG. 3.**
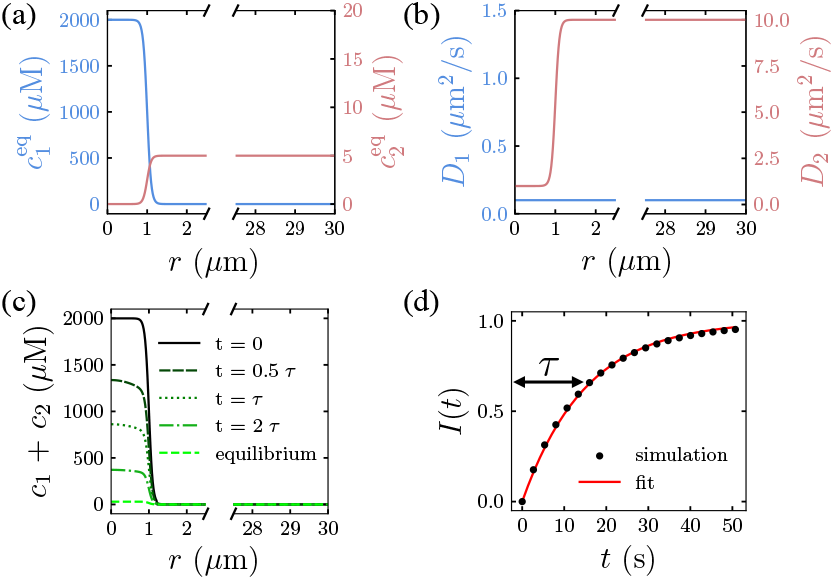
Representative simulation in a dilute-phase diffusion-limited scenario (parameters from top row in Table I). (a) Equilibrium concentration profiles and (b) diffusivity profiles for species 1 and 2. (c) Simulated radial concentration profiles of the bleached molecules for a few illustrative times. (d) Simulated brightness curve with exponential fit using nonlinear least squares [56]. Simulations were performed with radial step size *dr* = 20 nm over a system size of *r*_max_ = 30 *µ*m for 1000 recorded timepoints (the solver dynamically selects both the timestep and order for time integration).

We then solve Eqs. (1a) and (1b) numerically under spherical symmetry, using the initial and boundary conditions given by Eqs. (2) and (3), respectively, except that the boundary at *r* = +∞ is replaced by *r* = *r*_max_ for the finite size of the system. These coupled partial differential equations were solved using the *pdepe* function in MATLAB, which employs finite-difference spatial discretization with a variable-step, variable-order solver for time integration [55]. The numerical solutions for *c*_1_(*r, t*) and *c*_2_(*r, t*) (Fig. 3c) are used to compute a brightness curve in accordance with Eq. (4), which is subsequently fitted to extract the recovery timescale (Fig. 3d).

We show an example where the FRAP recovery is limited by dilute-phase diffusion in Fig. 3. Guided by Eq. (15), we choose physiological parameters of condensate systems [22, 57, 58] that lead to *τ* ≈ *τ*_dil_ (Table I). The numerically extracted relaxation time *τ* = 15.1 s of an *R* = 1 *μ*m droplet is indeed close to the theory prediction of *τ*_dil_ = 13.3 s. We repeat a similar procedure for various parameter sets in which the timescale of fluorescence recovery is limited by interfacial attachment/detachment, dense-phase diffusion of the low-mobility species, dense-phase diffusion of the high-mobility species, and attachment/detachment throughout the bulk of the condensate, totaling five cases. Simulated spatial fluorescence recovery profiles for each of these cases are shown in Fig. 4 with parameters listed in recovery (rows c and d) can readily be distinguished by the pronounced gradient present due to unbleached molecules gradually diffusing into the condensate and bleached ones diffusing out, whereas the remaining three cases all display a uniform recovery as the fast diffusion of molecules within the condensate quickly homogenizes the intensity profile. Details of the numerical implementation and fitting are provided in the Supplemental Material [51].

**FIG. 4.**
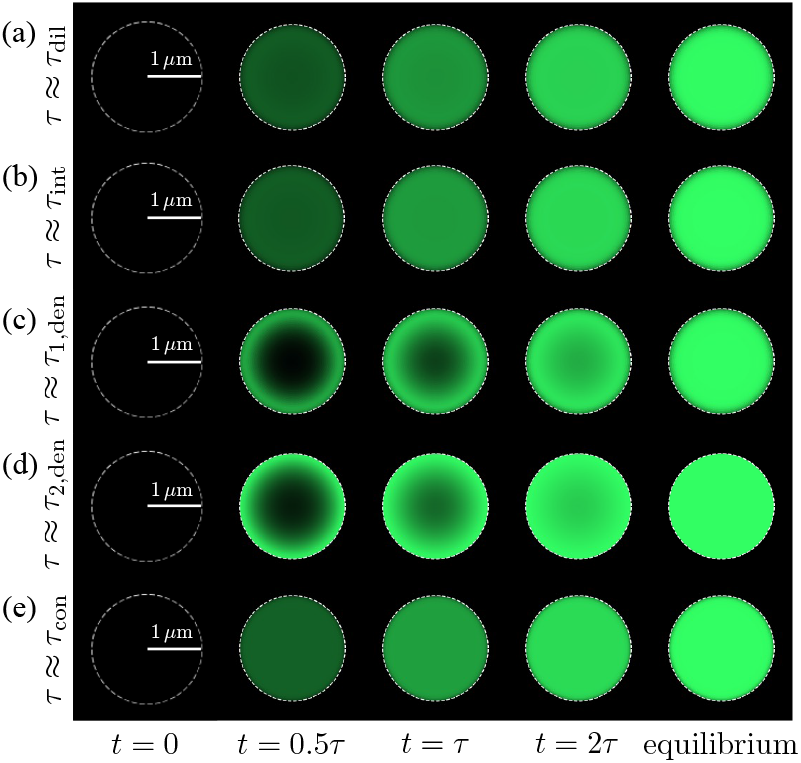
Simulated FRAP recovery profiles when fluorescence recovery is limited by (a) dilute-phase diffusion, (b) interfacial attachment/detachment, (c) dense-phase diffusion of species 1, (d) dense-phase diffusion of species 2, and (e) attachment/detachment throughout the bulk of the condensate. Green indicates fluorescent molecules and black indicates bleached molecules. Simulations were performed with radial step sizes 1*/*5 of the interface width *l*, and the system size was *r*_max_ = 30 *µ*m. 1000 timepoints were recorded over 4*τ* -long runtimes.

Based on Eqs. (5), (9), and (14), condensate recovery timescales are expected to follow different scaling laws in different rate-limiting cases. As shown in Fig. 5, the different timescales indeed scale differently with droplet size *in silico* as well. When diffusive processes are rate-limiting, the scaling law is quadratic; when the interfacial flux is rate-limiting, the scaling law is linear; and when the network attachment/detachment throughout the droplet is rate-limiting, the scaling law is independent of droplet size. We note that the theoretical curves in Fig. 5 are predictions from Eqs. (5), (9), and (14) with parameters in Table I, without any fitting to the simulated results. The deviations between theory and simulations in Figs. 5c and 5d reflect the presence of fast-decaying modes with non-negligible amplitudes in the diffusive processes [see Eq. (S40) in Supplemental Material]. Such deviations can be mitigated by fitting the simulated *I*(*t*) curves in the time range *t* ≥ 0.5*τ* to a function 1 − *A* exp( − *t*/*τ*), depicted as gray diamonds in Figs. 5c and 5d.

**TABLE 1.** Parameters for numerical simulations of five rate-limiting cases.

| Case | $c_{1,\text{den}}$<br>($\mu\text{M}$) | $c_{1,\text{dil}}$<br>($\mu\text{M}$) | $c_{2,\text{den}}$<br>($\mu\text{M}$) | $c_{2,\text{dil}}$<br>($\mu\text{M}$) | $D_{1,\text{den}}$<br>($\mu\text{m}^2/\text{s}$) | $D_{1,\text{dil}}^{\text{a}}$<br>($\mu\text{m}^2/\text{s}$) | $D_{2,\text{den}}$<br>($\mu\text{m}^2/\text{s}$) | $D_{2,\text{dil}}$<br>($\mu\text{m}^2/\text{s}$) | $l$<br>($\mu\text{m}$) | $k$<br>($\mu\text{M}^{-1}\text{s}^{-1}$) |
| --- | --- | --- | --- | --- | --- | --- | --- | --- | --- | --- |
| $\tau \approx \tau_{\text{dil}}$ | 2000 | 0 | 0 | 5 | 0.1 | 0.1 | 1 | 10 | 0.1 | 1 |
| $\tau \approx \tau_{\text{int}}$ | 1000 | 0 | 0 | 10 | 0.02 | 0.02 | 1 | 50 | 0.1 | 0.005 |
| $\tau \approx \tau_{1,\text{den}}$ | 1000 | 0 | 0 | 10 | 0.02 | 0.02 | 1 | 50 | 0.1 | 1 |
| $\tau \approx \tau_{2,\text{den}}$ | 1995 | 0 | 5 | 10 | $5 \times 10^{-5}$ | $5 \times 10^{-5}$ | 0.1 | 50 | 0.01 | 0.005 |
| $\tau \approx \tau_{\text{con}}$ | 980 | 0 | 20 | 10 | 0.02 | 0.02 | 1 | 50 | 0.05 | $1 \times 10^{-4}$ |

<sup>a</sup> Since there are no species 1 molecules in the dilute phase, we choose $D_{1,\text{dil}} = D_{1,\text{den}}$ to parameterize the diffusivity profile. This choice sets the diffusion coefficient of species 1 to be constant both inside the droplet and at its interface.

**FIG. 5.**
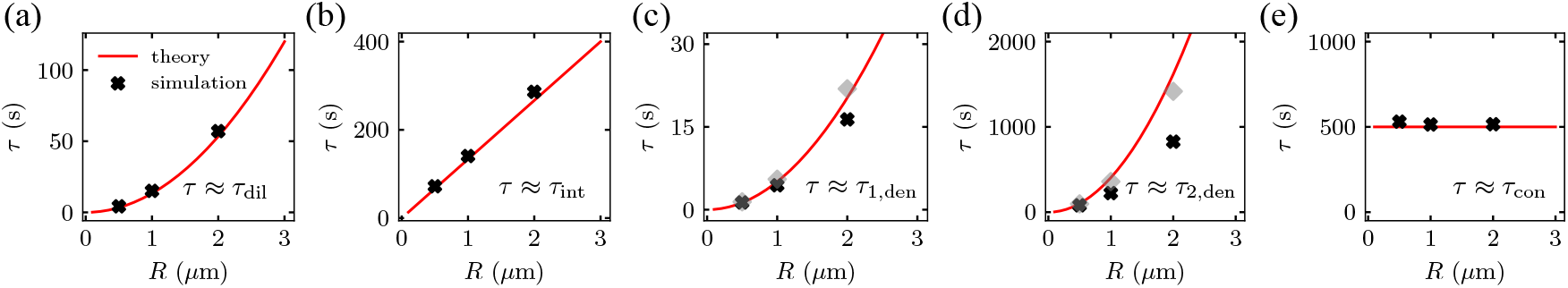
Theoretical and simulated scaling laws show good agreement in each rate-limiting case: (a) dilute-phase diffusion-limited timescale scales with *R*^2^, (b) interface-limited timescale scales with *R*, (c) dense-phase diffusion-limited timescale (species 1) scales with *R*^2^, (d) dense-phase diffusion-limited timescale (species 2) scales with *R*^2^, and (e) bulk conversion-limited timescale is independent of *R*. Red curves: theoretical predictions from Eqs. (5), (9), and (14) using parameters from Table I; black crosses: simulation results for droplets of radii *R* = 0.5 *µ*m, 1 *µ*m, and 2 *µ*m. Gray diamonds in (c) and (d) are from the same simulation results fit to the modified function 1 − *A* exp( − *t/τ* ) over a time range *t* ≥ 0.5*τ*, which disentangles the leading mode from the fast-decaying modes in diffusive processes.

Lastly, to provide a more complete picture, we present an example of a two-step recovery for a condensate with a significant high-mobility fraction (i.e., *c*_2,den_ ∼ *c*_1,den_). As discussed in the *Analytical derivations* subsection, when diffusion is faster than conversion, the droplet recovery consists of an initial fast mode due to the rapid replacement of bleached species 2 molecules by unbleached molecules diffusing in from the dilute phase, followed by a second slower mode due to the gradual detachment of bleached species 1 molecules from the network. As shown in Fig. S1b, the exchange dynamics proceeds in stages, giving rise to a double-exponential recovery.

### Application to a DNA nanostar system

Finally, we sought to employ our theory in an experimental system composed of DNA nanostars — a model system for investigating biomolecular condensation. Thanks to advances in DNA synthesis techniques, DNA nanostars offer highly programmable interactions: binding specificity and affinity can be tuned via the sequence and length of single-stranded overhangs, and valence via the number of arms. These features collectively enable a diverse range of phase behaviors [59–62]. Our DNA nanostars are composed of three arms of doublestranded DNA, each with a short tail of single-stranded DNA known as a “sticky end” due to its propensity to Watson-Crick base-pair with complementary strands (Fig. 6a) [63]. The sticky ends make these nanostars readily phase-separable, and micrometer-sized droplets can be seen with confocal microscopy (Figs. 6b and S2). Z-stack imaging also confirms that these droplets are mostly spherical (Fig. S2). Experiments were performed in a 40 mM Tris–Cl buffer solution at pH 7.9 containing 2.5 *μ*M DNA nanostars, 1 mM dithiothreitol (DTT), 27.5 mM magnesium acetate, and 2.5 mM nucleoside triphosphates (NTPs) at 37 ^*°*^C. Additional details about sequences and sample preparation are given in the Supplemental Material [51].

**FIG. 6.**
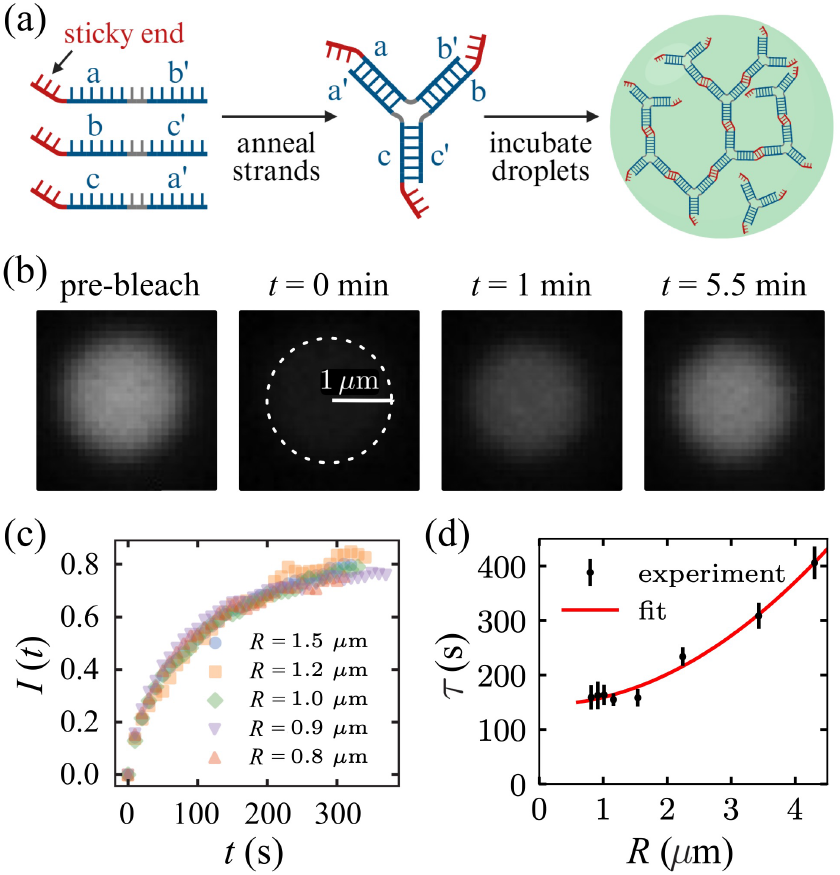
Experimental characterization of exchange dynamics in DNA nanostar condensates. (a) Nanostar schematic. The Y-shaped nanostar has 16-bp double-stranded arms (shown in blue), sticky ends with the palindromic sequence 5’- GCTAGC-3’ (shown in red), and 5’-TT-3’ domains between each of the three arms at the nanostar center (shown in gray) that confer angular flexibility between arms. The nanostars were first annealed in a low-salt solution, then added to a higher salt solution at 37 °C to incubate condensates, following the same protocol as in [63]. (b) Representative snapshots from a FRAP experiment of an *R* = 1 *µ*m nanostar droplet. (c) FRAP recovery curves for droplets of small sizes (*R* ≲ 1.5 *µ*m). (d) Exchange timescale versus droplet radius for all droplets, fitted with a shifted quadratic function of the form *τ* = *a* + *bR*^2^, where *a* = 145.0 *±* 5.6 s and *b* = 14.1 *±* 0.7 *µ*m^−2^s. Error bars on the data points represent 3*σ*. A replicate under the same experimental conditions is shown in Fig. S3, which confirms the same scaling behavior.

DNA nanostars form porous networks inside their condensates, with the volume fraction occupied by the nanostars on the order of 1% or less [61, 64]. This property makes them a prime system for observing the pore-mediated mode of recovery discussed above: nearby molecules may penetrate the droplet surface and diffuse freely within the droplet before attaching to the network. If this is the dominant recovery mechanism, we expect to observe conversion-limited recovery for small droplets, transitioning to diffusion-limited recovery for large droplets. Upon performing FRAP on nanostar droplets of varying sizes (Fig. 6b), we noticed that the recovery curves and hence the recovery timescales were nearly identical for droplets of small sizes (*R* ≲ 1.5 *μ*m), despite spanning nearly a twofold size range (Fig. 6c and 6d). This observation aligns with the scaling behavior of bulk conversion-limited recovery (*τ*_con_). If the recovery of these droplets were diffusion-limited, whether by dilute-phase diffusion or dense-phase diffusion (of either species), we would expect nearly a fourfold difference in exchange timescale. For larger droplets, we see a quadratic scaling that arises above the lower-limit plateau set by the bulk conversion-limited timescale (Fig. 6d). Upon fitting the data with the shifted quadratic function *τ* = *a* + *bR*^2^, we find the constants *a* and *b* are well-constrained: *a* = 145.0 *±* 5.6 s and *b* = 14.1 *±* 0.7 *μ*m^−2^s. The observed scalings were independently validated by a replicate experiment (Fig. S3).

As suggested by our theory, the plateau regime arises because the recovery is bulk conversion-limited, i.e., *τ*_con_ = 1/(*kc*_2,den_) = 145 s. In contrast to the detachment timescale of a nanostar from the network inferred here, the dissociation time of a single sticky end (6 bp) is typically much shorter, in the range *τ*_b_ = 0.01 – 10 s [65–67]. The difference can be explained by multivalency, which enhances the binding affinity [68, 69] and thereby extends the effective lifetime of a nanostar far beyond that of a single sticky end. Interestingly, the dissociation time of a DNA origami dimer held together by three 6bp stickers has been measured to be on the order of 100 s [70]. Given that physical parameters such as temperature and ionic strength can strongly impact the detachment timescales of both a single sticker and an entire nanostar [71], quantitative verification of the relevant timescales in our system will require future experiments, for example, via single-molecule tracking.

Nanostar droplets are porous, with a measured pore size comparable to the arm length. It has been reported that the partition coefficient for 70 kDa dextran (hydrodynamic radius 6 nm) is about 0.3 – 0.6 in such systems [64, 72]. Given that our nanostars have a hydrodynamic radius (5 – 7 nm) similar to that of dextran, we expect the unbound species of nanostars to partition in these nanostar droplets to a similar extent as the dextran. Assuming a partition coefficient of 0.5 and dilute-phase concentration of 1 *μ*M [63, 73], *c*_2,den_ ≈ 0.5 *c*_2,dil_ = 0.5 *μ*M, and the rate of nanostar attachment inside the condensate can be estimated as *k* = 1/(*ac*_2,den_) ≈ 0.014 *μ*M^−1^s^−1^. This rate appears to be much lower than the reported on-rate for nanostars in dilute solution, which ranges from 0.1 to 1 *μ*M^−1^s^−1^ [65, 74, 75]. The discrepancy likely arises because we have modeled the attachment flux as *kc*^eq^*c*_2_ in Eq. (1), implicitly assuming that every nanostar in the percolated network (species 1) can bind to freely-diffusing species 2 nanostars. However, many nanostars in the network may already be in a fully bound state or spatially occluded and thus unavailable for binding, leading to an apparent smaller value of the attachment rate *k*. Assuming nanostar attachment in the dense phase occurs with the same on-rate as in dilute solution, we can infer that more than 90% of nanostar sticky ends are not available for binding in these droplets.

The plateau in the *τ* versus *R* curve for small droplets also suggests that the interface-limited timescale *τ*_int_ is much longer than the bulk conversion-limited timescale *τ*_con_, i.e., *τ*_int_/*τ*_con_ = *Rc*_2,den_/(3*δ*_eff_*c*_2,dil_) ≫ 1. Nanostar condensates have a surface tension of *σ* ≈ 1*μ*N/m [61], which can be used to estimate the interfacial width as *l* ∼ (*k*_B_*T*/*σ*) ^½^ = 60 nm [64] and a corresponding effective width of *δ*_eff_ ≈ *l*/2 = 30 nm (see Supplemental Material [51]). Thus, even for the smallest measured droplet (*R* = 0.84 *μ*m), we have *τ*_int_/*τ*_con_ ≈ 5, indicating that *τ*_int_ is indeed much longer than *τ*_con_ for all droplets studied in these experiments. This also suggests that if no high-mobility species were present inside the nanostar droplets, the exchange timescale would be at least five times longer for these measured droplets, as predicted by Eq. (15).

Beyond the plateau regime, the negligible contribution of *τ*_int_ to the overall recovery time in Eq. (15) leads to a reduced expression for the recovery time:

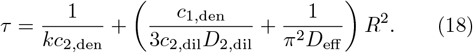

While the first term originated from *τ*_con_ sets the plateau value *a*, the second term from *τ*_dil_, *τ*_1,den_, and *τ*_2,den_ scales with *R*^2^ and sets the prefactor *b* = *c*_1,den_/(3*c*_2,dil_*D*_2,dil_) + 1/(*π*^2^*D*_eff_). Taking *c*_1,den_ = 200 *μ*M, *c*_2,dil_ = 1 *μ*M, and *D*_2,dil_ = 50 *μ*m^2^/s (from the Stokes-Einstein relation) [63, 73], we find that *τ*_dil_ ≪ *τ* for all droplets, suggesting that recovery is not limited by dilute-phase diffusion but by dense-phase diffusion. We then estimate *D*_eff_ ≈ 8 *×* 10^−3^*μ*m^2^/s from the fitted value of *b*, which sets upper bounds for *D*_1,den_ and *D*_2,den_: *D*_1,den_ ≤ 8 *×* 10^−3^*μ*m^2^/s and *D*_2,den_ ≤ 3 *μ*m^2^/s based on Eq. (8), assuming *c*_2,den_ = 0.5 *μ*M.

## DISCUSSION

A hallmark of biomolecular condensates is their dynamic exchange of material with their surroundings, a feature often crucial to their function. In this work, we developed a novel reaction-diffusion model to describe condensate exchange dynamics and explored how such dynamics can deviate from those of conventional liquid droplets due to the formation of a percolated network in the condensate. We found that in the presence of two mobility states, material exchange can be accelerated via a pore-mediated pathway in which molecules pass through the pores of the meshwork and attach/detach directly in the condensate interior. Notably, this pathway leads to a new regime where the exchange timescale becomes independent of condensate size, a prediction we confirmed using FRAP experiments on DNA nanostar condensates. In this study, we focused on the exchange dynamics of *in vitro* single-component condensates and approximated a condensate as a two-state system at equilibrium. However, molecules inside biological condensates, even in single-component systems, are likely to form a variable number of bonds rather than being simply fully bound or fully unbound. As a result, molecules inside condensates may exhibit a broad range of mobilities due to the multivalency of underlying interactions [49, 50]. Moreover, the relative abundances of different mobility states can evolve over time, for example, during condensate aging [76]. Reducing such systems to a two-state description can, in principle, give rise to non-Markovian effects that our minimal model does not capture. A natural next step would be to incorporate a distribution of mobility states into the model. Nevertheless, a concise two-state model can capture the essential physical principles underlying condensate exchange dynamics while remaining analytically tractable. Thus, our work serves as a first step toward addressing more complex dynamical problems in phase-separated systems.

The developed reaction-diffusion model can also be readily extended to describe multi-component systems. Condensates in living cells are complex assemblies of distinct proteins and nucleic acids, and much theoretical effort has been dedicated to understanding thermodynamic principles in multi-component systems with many coexisting phases [77–80]. However, less effort has been directed toward understanding the dynamics of such systems. Multi-component condensates generally employ a scaffold-client framework [30, 81]. While our model focused on the exchange dynamics of scaffold molecules, it can easily be generalized to describe client dynamics, where client molecules bind to/unbind from an equilibrated scaffold network, thus switching between low and high mobilities [82]. Since clients may bind not only to the percolated network (species 1) but also to unbound scaffold molecules (species 2), modeling client dynamics in this paradigm may require introducing a third species. In a similar spirit, the developed model can also be extended to predict the dynamics of condensates in the cell nucleus, such as transcriptional condensates and doublestrand break repair condensates, in which molecules diffusing within the condensate are also capable of binding to/unbinding from DNA [83–86].

The pore-mediated pathway involves molecules passing through the pores of the meshwork and attaching/detaching directly within the condensate. In which systems, and for which molecules, would this pathway be favored and experimentally measurable? Because diffusion within the connected network for bound molecules is generally much slower than for unbound molecules, the essential requirement for this pathway to dominate is that the mesh size of the percolated network be comparable or larger than the size of the molecule of interest, ensuring a sufficient concentration of high-mobility species. It has been reported that the mesh size (or correlation length) for *in vitro* reconstituted condensates of purified LAF-1 protein is about 6 nm [87], while for reconstituted coacervates composed of two oppositely charged intrinsically disordered proteins, histone H1 and prothymosin- *α*, it is 2.4 – 4.3 nm [88]. More generally, in a recent work, we established a quantitative relationship between mesh size and concentration, which suggests that condensates at physiologically relevant concentrations of 100 – 400 mg/ml exhibit mesh sizes decreasing from 8 to 3 nm [89]. Since protein sizes span the reported mesh size range, we expect the pore-mediated pathway to be relevant to the exchange dynamics of scaffolds in condensates with low dense-phase concentrations and to the exchange dynamics of clients more generally.

What is the biological implication of the pore-mediated pathway of condensate exchange dynamics? Although the bulk viscosity of condensates can be orders of magnitude higher than that of conventional oil droplets, small molecules can sneak through the pores of the meshwork and enter the condensate interior via this pore-mediated pathway, thereby bypassing slow diffusion caused by such high viscosity. The accelerated material exchange and diffusion enabled by this pathway are therefore expected to speed up biochemical reactions in metabolic condensates, as well as facilitate the response of condensates to changes in the intracellular environment more generally. If so, could the porosity of condensates be evolutionarily selected to promote more efficient dynamics?

Fascinating soft matter systems in their own right, biomolecular condensates are also increasingly implicated in cellular physiology and disease [90, 91]. We hope that our work will motivate further theoretical and experimental investigations into the complex dynamics in multicomponent, multi-state condensates, shedding light on their functional roles and paving the way for applications in condensate bioengineering.

## ACKNOWLEDGEMENTS

We are grateful to Ned Wingreen, Frank Jülicher, Alexander Grosberg, Jeremy Schmit, and Omar Saleh for insightful discussions and constructive feedback on the manuscript. R.K. and Y.Z. were supported by a startup fund at Johns Hopkins University. Y.Z. acknowledges support from the Sloan Foundation through a Sloan Research Fellowship. E.K. acknowledges support from Sloan Foundation grant 147170 to R.S. and NIH grant T32GM080189. R.S. acknowledges support from Sloan Foundation grant 147170, DOE BES award DESC0010426, NSF FET-2107246, ONR N00014-23-1-2868, and the Kent Gordon Croft Investment Management Faculty Scholar fund. This research was also supported in part by grant NSF PHY-2309135 to the Kavli Institute for Theoretical Physics (KITP).

