## Supplementary Material for "Dissecting Rate-Limiting Processes in Biomolecular Condensate Exchange Dynamics"

### Supplemental Material for "Dissecting Rate-Limiting Processes in Biomolecular Condensate Exchange Dynamics"

(Dated: December 16, 2025)

### I. Supplementary Analytical Derivations

#### a. Derivation of the dilute-phase diffusion-limited timescale in Equation (5a)

Imposing spherical symmetry on Eqs. (1a) and (1b) in the main text, we start with the following governing equations:

$$\frac{\partial c_1(r,t)}{\partial t} = \frac{1}{r^2}\frac{\partial}{\partial r}\left\{r^2 D_1(r)\left[\frac{\partial c_1(r,t)}{\partial r} - \frac{c_1(r,t)}{c_1^{\text{eq}}(r)}\frac{dc_1^{\text{eq}}(r)}{dr}\right]\right\} + k\left[c_1^{\text{eq}}(r)c_2(r,t) - c_2^{\text{eq}}(r)c_1(r,t)\right], \quad \text{(S1a)}$$

$$\frac{\partial c_2(r,t)}{\partial t} = \frac{1}{r^2}\frac{\partial}{\partial r}\left\{r^2 D_2(r)\left[\frac{\partial c_2(r,t)}{\partial r} - \frac{c_2(r,t)}{c_2^{\text{eq}}(r)}\frac{dc_2^{\text{eq}}(r)}{dr}\right]\right\} - k\left[c_1^{\text{eq}}(r)c_2(r,t) - c_2^{\text{eq}}(r)c_1(r,t)\right]. \quad \text{(S1b)}$$

Outside the condensate, Eq. (S1) reduces to

$$\frac{\partial c_2(r,t)}{\partial t} = D_{2,\text{dil}}\frac{1}{r^2}\frac{\partial}{\partial r}\left[r^2\frac{\partial c_2(r,t)}{\partial r}\right], \quad \text{(S2)}$$

since $c_1(r,t) = 0$, $c_1^{\text{eq}}(r) = 0$, $dc_2^{\text{eq}}(r)/dr = 0$, and $D_2(r) = D_{2,\text{dil}}$.

In the dilute-phase diffusion-limited regime, both the diffusion timescale inside the condensate and the conversion timescale between species 1 and 2 are faster than the recovery timescale. Therefore, the bleached concentrations $c_1(r,t)$ and $c_2(r,t)$ can be viewed as equilibrated within the condensate:

$$c_i(r,t) \approx \alpha(t)c_i^{\text{eq}}(r), \quad \text{(S3)}$$

with time-dependent component $\alpha(t)$ and $i = 1, 2$. Adding Eqs. (S1a) and (S1b) and then applying the divergence theorem to the resulting expression yields

$$\frac{d}{dt}\int_0^{R^+}\left[c_1(r,t) + c_2(r,t)\right]4\pi r^2 dr = 4\pi r^2 D_2(r)\frac{\partial}{\partial r}c_2(r,t)\bigg|_{r=R^+}, \quad \text{(S4)}$$

where $R^+$ denotes a location just outside the condensate. Physically, this means that the rate of change in the number of bleached molecules within a volume is equal to the net flux through its surface. Plugging the approximate forms of $c_1(r,t)$ and $c_2(r,t)$ [Eq. (S3)] into the above equation, we get

$$\frac{4\pi R^3\left(c_{1,\text{den}} + c_{2,\text{den}}\right)}{3}\frac{d\alpha(t)}{dt} = 4\pi R^2 D_{2,\text{dil}}\frac{\partial}{\partial r}c_2(r,t)\bigg|_{r=R^+}, \quad \text{(S5)}$$

where we have assumed a narrow interface width so that the integral on the left-hand side of Eq. (S4) is dominated by contributions from the bulk of the condensate.

Local equilibrium at the droplet interface relates the bleached concentrations inside and outside the droplet:

$$c_2(r = R^+, t) = \alpha(t) c_2^{\text{eq}}(r = R^+) = \alpha(t) c_{2,\text{dil}}, \tag{S6}$$

and we expect there to be no bleached molecules infinitely far from the droplet:

$$c_2(r = +\infty, t) = 0. \tag{S7}$$

Taken together, these two boundary conditions can be applied to Eq. (S2) to obtain a steady-state solution:

$$c_2^{\text{ss}}(r, t) = \alpha(t) c_{2,\text{dil}} \frac{R}{r}, \tag{S8}$$

which is a good approximation to the solution of Eq. (S2) under the assumption that the rate of change of $\alpha(t)$ is much slower than the convergence rate of the solution ($\sim D_{2,\text{dil}}/R^2$) to its steady-state form, $c_2^{\text{ss}}(r, t)$. Substituting the steady-state solution of $c_2(r, t)$ into Eq. (S5) leads to

$$\frac{R^2 \left(c_{1,\text{den}} + c_{2,\text{den}}\right)}{3} \frac{d\alpha(t)}{dt} = -D_{2,\text{dil}} c_{2,\text{dil}} \alpha(t). \tag{S9}$$

Therefore,

$$\alpha(t) = e^{-\frac{t}{\tau_{\text{dil}}}}, \tag{S10}$$

where

$$\tau_{\text{dil}} = \frac{R^2 c_{1,\text{den}}}{3 D_{2,\text{dil}} c_{2,\text{dil}}}, \tag{S11}$$

assuming $c_{2,\text{den}} \ll c_{1,\text{den}}$. We note that the rate of change of $\alpha(t)$, $1/\tau_{\text{dil}}$, is indeed much slower than $D_{2,\text{dil}}/R^2$, thereby validating the solution in Eq. (S8). Finally, the normalized brightness curve $I(t)$ for the fraction of unbleached molecules inside the droplet is

$$I(t) = 1 - \alpha(t) = 1 - e^{-\frac{t}{\tau_{\text{dil}}}}, \tag{S12}$$

identifying $\tau_{\text{dil}}$ as the dilute-phase diffusion-limited timescale.

##### b. Derivation of the interface conversion-limited timescale in Equation (5b)

To derive the interface conversion-limited timescale, we first apply the divergence theorem to Eqs. (S1a) and (S1b), leading to:

$$\frac{d}{dt} \int_0^\infty c_1(r, t) 4\pi r^2 dr = \int_0^\infty k \left[c_1^{\text{eq}}(r) c_2(r, t) - c_2^{\text{eq}}(r) c_1(r, t)\right] 4\pi r^2 dr, \tag{S13a}$$

$$\frac{d}{dt}\int_0^\infty c_2(r,t)4\pi r^2 dr = \int_0^\infty k\left[c_2^{\mathrm{eq}}(r)c_1(r,t) - c_1^{\mathrm{eq}}(r)c_2(r,t)\right]4\pi r^2 dr. \tag{S13b}$$

We note that the diffusion terms in Eqs. (S1a) and (S1b) drop out because of the no-flux boundary conditions from Eq. (3) in the main text. Since diffusion is fast in this limit, the bleached molecules of each species can be considered equilibrated with respect to their own equilibrium concentration profiles, i.e., Eqs. (S13a) and (S13b) have separable solutions of the form

$$c_i(r,t) \approx \alpha_i(t)c_i^{\mathrm{eq}}(r), \tag{S14}$$

with time-dependent component $\alpha_i(t)$ and $i = 1,2$. Strictly speaking, within the conversion-limited timescale $\tau$, species 2 molecules can diffuse only over a distance $d \sim \sqrt{D_{\mathrm{dil}}\tau}$. Therefore, $c_2(r,t) \approx \alpha_2(t)c_2^{\mathrm{eq}}(r)$ in Eq. (S14) is valid only for a system of size $L \ll d$. Since $\tau \gg \tau_{\mathrm{dil}}$ by construction, $d \sim \sqrt{D_{\mathrm{dil}}\tau} \gg \sqrt{D_{\mathrm{dil}}\tau_{\mathrm{dil}}} \sim R\sqrt{c_{1,\mathrm{den}}/c_{2,\mathrm{dil}}}$, and a proper choice of $L$ can be identified as $L = R\sqrt{c_{1,\mathrm{den}}/c_{2,\mathrm{dil}}}$. For a system of larger size, $\alpha_2$ becomes spatially dependent. However, because the bleached concentration of species 2 in the dense phase quickly decreases due to diffusion into the dilute phase, we can estimate $\alpha_2(t) \le c_{2,\mathrm{den}}/[c_{2,\mathrm{den}} + c_{2,\mathrm{dil}}(L^3/R^3 - 1)] \ll 1$ at time $t$ longer than the relevant diffusion-limited timescales. Therefore, $\alpha_2(t) \approx 0$ and the approximation $c_2(r,t) \approx \alpha_2(t)c_2^{\mathrm{eq}}(r)$ is still valid for larger systems.

Substituting the approximate forms of $c_1(r,t)$ and $c_2(r,t)$ in Eq. (S14) into Eqs. (S13a) and (S13b) reduces the problem to a system of two simple ordinary differential equations:

$$\frac{d\alpha_1(t)}{dt} = k\frac{a}{N_1}\left[\alpha_2(t) - \alpha_1(t)\right], \tag{S15a}$$

$$\frac{d\alpha_2(t)}{dt} = k\frac{a}{N_2}\left[\alpha_1(t) - \alpha_2(t)\right], \tag{S15b}$$

where

$$a = \int_0^\infty c_1^{\mathrm{eq}}(r)c_2^{\mathrm{eq}}(r)4\pi r^2 dr, \tag{S16a}$$

$$N_i = \int_0^\infty c_i^{\mathrm{eq}}(r)4\pi r^2 dr, \tag{S16b}$$

with $i = 1,2$ and $N_i$ denoting the total number of species $i$ molecules. Eqs. (S15a) and (S15b) can be solved analytically:

$$\alpha_i(t) = (f_i - f_b)e^{-\frac{t}{\tau}} + f_b, \tag{S17}$$

where the constant $f_i$ denotes the fraction of molecules bleached in species $i$, and $f_b$ is the fraction of total bleached molecules. We identify the timescale for fluorescence recovery in the limit of slow conversion as $\tau = N_1 N_2 / [ka(N_1 + N_2)]$, which is explicitly:

$$\tau = \frac{\int_0^\infty c_1^{\text{eq}}(r) r^2 dr \int_0^\infty c_2^{\text{eq}}(r) r^2 dr}{k \int_0^\infty c_1^{\text{eq}}(r) c_2^{\text{eq}}(r) r^2 dr \int_0^\infty \left[c_1^{\text{eq}}(r) + c_2^{\text{eq}}(r)\right] r^2 dr}. \tag{S18}$$

In the limit where the dilute phase contains many more molecules than the bleached droplet (as is usually the case), Eq. (S18) simplifies to

$$\tau = \frac{\int_0^\infty c_1^{\text{eq}}(r) r^2 dr}{k \int_0^\infty c_1^{\text{eq}}(r) c_2^{\text{eq}}(r) r^2 dr}, \tag{S19}$$

and $f_1 = 1$ and $f_2 = f_b = 0$, leading to

$$I(t) = 1 - \frac{c_{1,\text{den}}}{c_{1,\text{den}} + c_{2,\text{den}}} \alpha_1(t) = 1 - \frac{c_{1,\text{den}}}{c_{1,\text{den}} + c_{2,\text{den}}} e^{-\frac{t}{\tau}}. \tag{S20}$$

In the absence of species 2 molecules in the bulk of the droplet, i.e., $c_{2,\text{den}} = 0$, attachment and detachment can only occur at the droplet interface. In this case, the integral form of $\tau$ in Eq. (S19) reduces to the interface-limited timescale:

$$\tau_{\text{int}} = \frac{R}{3kc_{2,\text{dil}}\delta_{\text{eff}}}, \tag{S21}$$

where

$$\delta_{\text{eff}} \equiv \frac{\int_0^\infty c_1^{\text{eq}}(r) c_2^{\text{eq}}(r) r^2 dr}{c_{1,\text{den}} c_{2,\text{dil}} R^2}, \tag{S22}$$

and $I(t) = 1 - e^{-t/\tau_{\text{int}}}$. We note that the integrand in the numerator of Eq. (S22) is non-zero only at the droplet interface, which gives rise to a factor of $R^2$ that cancels the $R^2$ in the denominator. For given concentration profiles:

$$c_1^{\text{eq}}(r) = \frac{c_{1,\text{den}}}{2} - \frac{c_{1,\text{den}}}{2} \tanh\left(\frac{r-R}{l}\right), \tag{S23a}$$

$$c_2^{\text{eq}}(r) = \frac{c_{2,\text{dil}}}{2} + \frac{c_{2,\text{dil}}}{2} \tanh\left(\frac{r-R}{l}\right), \tag{S23b}$$

our numerical results suggest $\delta_{\text{eff}} \approx l/2$.

##### c. Derivation of the dense-phase low-mobility species diffusion-limited timescale in Equation (5c)

In the limit where dense-phase diffusion is much slower than dilute-phase diffusion and network attachment/detachment, the bleached concentrations of species 1 and 2 can be considered locally equilibrated with each other:

$$c_i(r,t) \approx \alpha(r,t) c_i^{\mathrm{eq}}(r). \tag{S24}$$

To derive Eq. (6) in the main text, we first substitute Eq. (S24) into Eqs. (S1a) and (S1b), and rewrite the diffusion terms as:

$$\frac{\partial c_1(r,t)}{\partial t} = \frac{1}{r^2}\frac{\partial}{\partial r}\left\{ r^2 D_1(r) c_1(r,t) \left[ \frac{\partial \ln c_1(r,t)}{\partial r} - \frac{d \ln c_1^{\mathrm{eq}}(r)}{dr} \right] \right\}, \tag{S25a}$$

$$\frac{\partial c_2(r,t)}{\partial t} = \frac{1}{r^2}\frac{\partial}{\partial r}\left\{ r^2 D_2(r) c_2(r,t) \left[ \frac{\partial \ln c_2(r,t)}{\partial r} - \frac{d \ln c_2^{\mathrm{eq}}(r)}{dr} \right] \right\}. \tag{S25b}$$

We note that

$$\ln c_1(r,t) - \ln c_1^{\mathrm{eq}}(r) = \ln c_2(r,t) - \ln c_2^{\mathrm{eq}}(r) = \ln \alpha(r,t) = \ln c(r,t) - \ln c^{\mathrm{eq}}(r), \tag{S26}$$

where

$$c(r,t) = c_1(r,t) + c_2(r,t), \tag{S27a}$$

$$c^{\mathrm{eq}}(r) = c_1^{\mathrm{eq}}(r) + c_2^{\mathrm{eq}}(r). \tag{S27b}$$

Adding Eqs. (S25a) and (S25b) then leads to

$$\frac{\partial c(r,t)}{\partial t} = \frac{1}{r^2}\frac{\partial}{\partial r}\left\{ r^2 \left[ D_1(r) c_1(r,t) + D_2(r) c_2(r,t) \right] \left[ \frac{\partial \ln c(r,t)}{\partial r} - \frac{d \ln c^{\mathrm{eq}}(r)}{dr} \right] \right\}, \tag{S28}$$

which is equivalent to

$$\frac{\partial c(r,t)}{\partial t} = \frac{1}{r^2}\frac{\partial}{\partial r}\left\{ r^2 D(r) \left[ \frac{\partial c(r,t)}{\partial r} - \frac{c(r,t)}{c^{\mathrm{eq}}(r)} \frac{d c^{\mathrm{eq}}(r)}{dr} \right] \right\}, \tag{S29}$$

where

$$D(r) = \frac{D_1(r) c_1^{\mathrm{eq}}(r) + D_2(r) c_2^{\mathrm{eq}}(r)}{c^{\mathrm{eq}}(r)}. \tag{S30}$$

In the dense-phase diffusion-limited regime, we solve the effective diffusion equation in Eq. (S29) subject to the following boundary conditions:

$$c(R,t) = 0, \tag{S31a}$$

$$\left.\frac{\partial c(r,t)}{\partial r}\right|_{r=0} = 0, \tag{S31b}$$

and initial condition:

$$c(r,0) = c^{\mathrm{eq}}(r), \quad r < R. \tag{S32}$$

Eq. (S31a) reflects that molecules are immediately absorbed into the dilute phase upon reaching the droplet surface, and Eq. (S31b) reflects that there is no flux of molecules at the center of the droplet. If we further consider a narrow interface width, Eq. (S29) reduces to

$$\frac{\partial c(r,t)}{\partial t} = D_{\mathrm{den}} \left( \frac{\partial^2}{\partial r^2} + \frac{2}{r}\frac{\partial}{\partial r} \right) c(r,t), \quad r < R, \tag{S33}$$

where

$$D_{\mathrm{den}} = \frac{D_{1,\mathrm{den}} c_{1,\mathrm{den}} + D_{2,\mathrm{den}} c_{2,\mathrm{den}}}{c_{1,\mathrm{den}} + c_{2,\mathrm{den}}} \approx D_{1,\mathrm{den}} + D_{2,\mathrm{den}} \frac{c_{2,\mathrm{den}}}{c_{1,\mathrm{den}}}, \tag{S34}$$

assuming $c_{1,\mathrm{den}} \gg c_{2,\mathrm{den}}$.

Equation (S33) can be solved using the method of separation of variables. Letting $c(r,t) = A(r)B(t)$, with spatial and time dependence given by functions $A(r)$ and $B(t)$, respectively, we can rewrite Eq. (S33) as

$$\frac{1}{D_{\mathrm{den}} B(t)} \frac{dB(t)}{dt} = -\lambda = \frac{1}{A(r)} \left( \frac{d^2 A(r)}{dr^2} + \frac{2}{r}\frac{dA(r)}{dr} \right). \tag{S35}$$

Since the left-hand side in Eq. (S35) only depends on $t$ and the right-hand side only depends on $r$, both sides must be equal to a constant $-\lambda$, where $\lambda > 0$ such that the solution decays over time. Solving for $A(r)$ and $B(t)$ gives the general solution:

$$A(r) = \frac{A_0 \sin(\sqrt{\lambda} r)}{r} + \frac{A_1 \cos(\sqrt{\lambda} r)}{r}, \tag{S36a}$$

$$B(t) = B_0 e^{-\lambda D_{\mathrm{den}} t}, \tag{S36b}$$

where $A_0$, $A_1$, and $B_0$ are constants. In order for the solution to be finite at the origin, $A_1$ must be zero. Then, applying the boundary condition in Eq. (S31a) to Eq. (S36a) gives the eigenvalues of $\lambda$ as $\lambda = n^2\pi^2/R^2$, where $n \in \{\pm 1, \pm 2, \ldots\}$. By linearity of the diffusion equation, the full solution to Eq. (S33) is the following superposition:

$$c(r,t) = \sum_{n=1}^{\infty} \frac{A_n}{r} \sin\left(\frac{n\pi r}{R}\right) \exp\left(-\frac{n^2\pi^2}{R^2} D_{\mathrm{den}} t\right), \tag{S37}$$

with the initial condition $c(r,0) = c_{\mathrm{den}}$ represented by its Fourier series as

$$c_{\mathrm{den}} = \sum_{n=1}^{\infty} \frac{A_n}{r} \sin\left(\frac{n\pi r}{R}\right), \quad r < R. \tag{S38}$$

Combined with Eq. (S38), the orthogonality condition for spherically symmetric basis functions in three dimensions determines the amplitudes $A_n$:

$$A_n = \frac{2c_{\text{den}}}{R} \int_0^R \sin\left(\frac{n\pi r}{R}\right) r dr = -\frac{2c_{\text{den}} R \cos(\pi n)}{\pi n}. \tag{S39}$$

The terms in the solution in Eq. (S37) decay according to $e^{-t/\tau_n}$, where $\tau_n = R^2/(n^2\pi^2 D_{\text{den}})$.

Having obtained $c(r,t)$, we can write the full solution for the brightness recovery curve:

$$I(t) = 1 - \frac{\int_0^R c(r,t) r^2 dr}{\int_0^R c_{\text{den}} r^2 dr} = 1 - \frac{6}{\pi^2} \sum_{n=1}^{\infty} \frac{1}{n^2} e^{-t/\tau_n}. \tag{S40}$$

This solution obeys the expected limits that $I(t=0) = 0$ and $I(t \to \infty) = 1$. Since the timescales $\tau_n$ decrease rapidly with increasing $n$, the long-term behavior of this solution is well-characterized by the $n = 1$ timescale:

$$\tau_{\text{den}} = \frac{R^2}{\pi^2 D_{\text{den}}}. \tag{S41}$$

The above derivation is general for dense-phase diffusion-limited recovery, but since we are in the dense-phase low-mobility species diffusion-limited regime, diffusion of species 2 does not contribute to the recovery, i.e., $D_{\text{den}} \approx D_{1,\text{den}}$ by construction. Therefore, the timescale associated with dense-phase diffusion of species 1 is

$$\tau_{1,\text{den}} = \frac{R^2}{\pi^2 D_{1,\text{den}}}. \tag{S42}$$

##### d. Derivation of the dense-phase high-mobility species diffusion-limited timescale in Equation (9)

The dense-phase high-mobility species diffusion-limited timescale can be derived following the same derivation as in the previous section. In this regime, however, diffusion of species 1 does not contribute to the recovery, i.e., $D_{\text{den}} \approx D_{2,\text{den}} c_{1,\text{den}}/c_{2,\text{den}}$ by construction [1, 2]. Therefore, the timescale associated with dense-phase diffusion of species 2 is

$$\tau_{2,\text{den}} = \frac{R^2 c_{1,\text{den}}}{\pi^2 D_{2,\text{den}} c_{2,\text{den}}}. \tag{S43}$$

##### e. Derivation of the bulk conversion-limited timescale in Equation (14)

The derivation in the interface conversion-limited subsection generally applies to the bulk conversion-limited regime as well. The main difference is the final simplifying assumption: here,

$c_{2,\mathrm{den}} \neq 0$, so attachment/detachment not only occurs at the interface of the droplet but also throughout the bulk of the condensate. When the bulk conversion dominates the recovery, Eq. (S19) reduces to the bulk conversion-limited timescale:

$$\tau_{\mathrm{con}} = \frac{1}{k c_{2,\mathrm{den}}}. \tag{S44}$$

#### II. Numerical Implementation and Fitting

##### a. Extracting the exchange timescale $\tau$

Using *SciPy*'s optimized curve fitting function, we fit both the numerical and experimental brightness curves with a single exponential function $I_{\mathrm{fit}}(t) = 1 - e^{-t/\tau}$. The trust region reflective method is used to minimize the sum of squared residuals [3]. While the total recovery is constrained to 1, we note that small numerical deviations of this fit may be partly due to the finite size of the simulated dilute phase, the multiple relaxation modes inherent in the dense-phase diffusion-limited cases (both species), or the nonzero contributions of secondary recovery processes. Accuracy of the fit with respect to the theoretical curves can be improved in the dense-phase diffusion limited cases (both species) by instead fitting only the $t \gtrsim 0.5\tau$ region with the function $1 - Ae^{-t/\tau}$. In experiments, another source of error may be a small immobile fraction that prevents total recovery to 1. As for fitting the recovery timescale versus droplet size, the same methods are used to extract parameters $a$ and $b$ in the shifted quadratic function $\tau = a + bR^2$.

##### b. Double exponential recovery

Throughout the main text, we have assumed that it is energetically favorable for molecules within the droplet to attach to the network rather than persist in a detached state, implying $c_{2,\mathrm{den}} \ll c_{1,\mathrm{den}}$ as determined by their Boltzmann factors. However, in certain situations, the high-mobility species can constitute a significant fraction of the dense phase, i.e., $c_{2,\mathrm{den}} \sim c_{1,\mathrm{den}}$. Following the discussion of this case in the main text, if conversion is slower than diffusion, the bleached molecules of species 2 inside the droplet will quickly be replaced by unbleached molecules diffusing in from the dilute phase, whereas the bleached molecules of species 1 can only be replaced through slow exchange with unbleached molecules of species 2 at the droplet interface and bulk, leading to a two-step recovery.

We show such an example in Fig. S1, where the small high-mobility fraction case ($c_{2,\mathrm{den}} \ll c_{1,\mathrm{den}}$) follows a single exponential recovery (Fig. S1a), while the large high-mobility fraction case ($c_{2,\mathrm{den}} \sim c_{1,\mathrm{den}}$) follows a double exponential recovery (Fig. S1b). Parameters for the two cases are listed in the top and bottom rows of Table S1, respectively. In both cases, we choose parameters such that the network attachment/detachment timescale in the droplet bulk is faster than that at the interface, but much slower than the dense- and dilute-phase diffusion timescales. For the small high-mobility fraction case (Fig. S1a), following Eq. (15) in the main text, the bulk conversion-limited timescale is expected to dominate the long-term recovery. The numerically extracted timescale $\tau = 532.3\,\mathrm{s}$ is indeed close to the theoretical prediction of $\tau_{\mathrm{con}} = 500\,\mathrm{s}$ for this case. For the large high-mobility fraction case (Fig. S1b), we follow the same fitting procedure described in the previous section but with a double exponential function of the form $A(1 - e^{-t/\tau_1}) + (1 - A)(1 - e^{-t/\tau_2})$. The fast recovery mode has an amplitude $A = 0.25$ and a timescale $\tau_1 = 0.2\,\mathrm{s}$, while the slow recovery mode has an amplitude $1 - A = 0.75$ and a timescale $\tau_2 = 47.8\,\mathrm{s}$. Physically, since the initial fast recovery is due to bleached species 2 molecules diffusing out of the dense phase and being replaced by unbleached species 2 molecules diffusing in from the dilute phase, we expect its associated timescale to match the diffusion-limited timescale for species 2: $R^2/(\pi^2 D_{2,\mathrm{den}}) + c_{2,\mathrm{den}} R^2/(3c_{2,\mathrm{dil}} D_{2,\mathrm{dil}}) = 0.27\,\mathrm{s}$. This expression is essentially a sum of the timescales in Eqs. (5a) and (5c), with parameters of species 1 replaced by those of species 2. The fast recovery mode accounts for $c_{2,\mathrm{den}}/(c_{1,\mathrm{den}} + c_{2,\mathrm{den}}) = 25\%$ of the total recovery, corresponding to the complete replacement of bleached high-mobility molecules in the droplet. The remaining 75% of the recovery is driven by the slow network detachment of bleached species 1 molecules and the subsequent attachment of unbleached species 2 molecules throughout the droplet bulk. The associated timescale of this slow recovery mode (47.8 s) is indeed close to the expected bulk conversion-limited timescale $\tau_{\mathrm{con}} = 40\,\mathrm{s}$.

TABLE S1. Parameters for numerical simulations of small (2%) and large (25%) high-mobility fraction cases. The simulated droplet radius is fixed at $R = 1\,\mu\mathrm{m}$ in both cases.

| Case | $c_{1,\mathrm{den}}$ ($\mu$M) | $c_{1,\mathrm{dil}}$ ($\mu$M) | $c_{2,\mathrm{den}}$ ($\mu$M) | $c_{2,\mathrm{dil}}$ ($\mu$M) | $D_{1,\mathrm{den}}$ ($\mu\mathrm{m}^2/\mathrm{s}$) | $D_{1,\mathrm{dil}}$ ($\mu\mathrm{m}^2/\mathrm{s}$) | $D_{2,\mathrm{den}}$ ($\mu\mathrm{m}^2/\mathrm{s}$) | $D_{2,\mathrm{dil}}$ ($\mu\mathrm{m}^2/\mathrm{s}$) | $l$ ($\mu$m) | $k$ ($\mu\mathrm{M}^{-1}\mathrm{s}^{-1}$) |
|---|---|---|---|---|---|---|---|---|---|---|
| small high-mobility fraction | 980 | 0 | 20 | 10 | 0.02 | 0.02 | 1 | 50 | 0.1 | $1 \times 10^{-4}$ |
| large high-mobility fraction | 750 | 0 | 250 | 10 | 0.02 | 0.02 | 1 | 50 | 0.1 | $1 \times 10^{-4}$ |

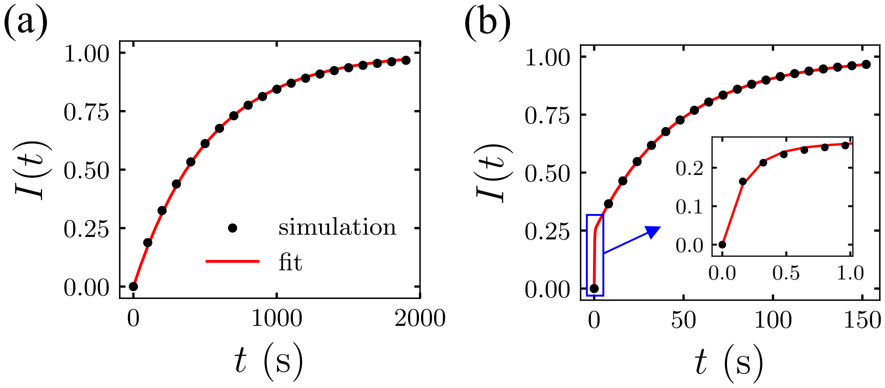


FIG. S1. Brightness curves illustrate (a) single exponential recovery for a small high-mobility fraction case and (b) double exponential recovery for a large high-mobility fraction case. Inset reveals single exponential behavior at early times.

#### III. DNA Nanostar Experimental Methods

The following methods are adapted from previous literature [4].

##### a. Materials and reagents

DNA strands were purchased from Integrated DNA Technologies (Coralville, IA). The fluorophore-containing strand was purified using high-performance liquid chromatography (HPLC), and all other strands were purified using a proprietary desalting technique from Integrated DNA Technologies. Capillary tubes and glass slides were purchased from VitroCom (Mountain Lakes, NJ) and VWR (Radnor, PA), respectively. Nucleoside triphosphates (NTPs) and dithiothreitol (DTT) were purchased from Thermo Fisher Scientific (Waltham, MA). Tris base, HCl, and magnesium acetate were purchased from Sigma-Aldrich (Allentown, PA).

##### b. Nanostar structure formation

DNA strands (Table S2) were added to a buffer solution (20 mM Tris-HCl at pH 7.9 and 1.33 mM magnesium acetate) to yield a final concentration of 20 $\mu$M DNA NS, of which 10% was fluorescently labeled with Atto647 fluorophore. The annealing protocol consisted of first raising the temperature to 90 °C, holding for 5 minutes, then decreasing 1 °C/min until reaching 20 °C. NS solutions were subsequently either maintained at 4 °C and used shortly thereafter, or stored at −20 °C for later

TABLE S2. DNA sequences used to form nanostar (NS) structures [5].

| Name | Sequence |
|---|---|
| YS1-1 | GCTAGCCAGTGAGGACGGAAGTTTGTCGTAGCATCGCACC |
| YS1-2 | GCTAGCCAACCACGCCTGTCCATTACTTCCGTCCTCACTG |
| YS1-3 | GCTAGCGGTGCGATGCTACGACTTTGGACAGGCGTGGTTG |
| YS1-1-Atto647 | /5Atto647NN/GCTAGCCAGTGAGGACGGAAGTTTGTCGTAGCATCGCACC |

use within a maximum of two freeze-thaw cycles.

##### c. Nanostar droplet formation

The preformed NSs were added to a solution of 27.5 mM magnesium acetate, 40 mM Tris-Cl pH 7.9, 1 mM DTT, and 2.5 mM NTPs, yielding a final NS concentration of 2.5 $\mu$M. Immediately upon addition of NS, the solution was drawn up into a capillary tube, sealed, and glued onto a glass slide using epoxy. The slide was then placed inside a microscope equipped with an incubation chamber set to 37 °C to form droplets.

##### d. Whole-droplet FRAP experiment

FRAP experiments were performed using an Olympus/Evident FV4000 laser scanning confocal microscope with a 60x-magnification, 1.4-numerical aperture oil-immersion objective, temperature incubation chamber set to 37 °C, and a 640 nm laser. Optical detection was fully spectrally determined (no emission filters) using two blue-shifted spectral detectors, two red-shifted spectral detectors, and one transmission photomultiplier tube. The incubation chamber was preincubated at 37 °C for at least 1 hour before placing samples inside. After placing the glass slide inside the chamber, the sample was left to equilibrate for another 30 minutes. To acquire z-stack images, the pinhole of the microscope was set to 1 Airy Unit (AU) (0.56 $\mu$m). Z-stack image sections were spaced apart by 0.5 AU. A representative z-stack is shown in Figure S2.

For FRAP of whole droplets, the pinhole of the microscope was set to its maximum setting of 3.05 AU. Immediately after taking the first image of the droplet, a region slightly larger than the droplet of interest was bleached using a 640 nm laser for the Atto647N dye. After bleaching, the droplet was monitored and imaged every 10 seconds to acquire time-lapse snapshots for subsequent recovery analysis.

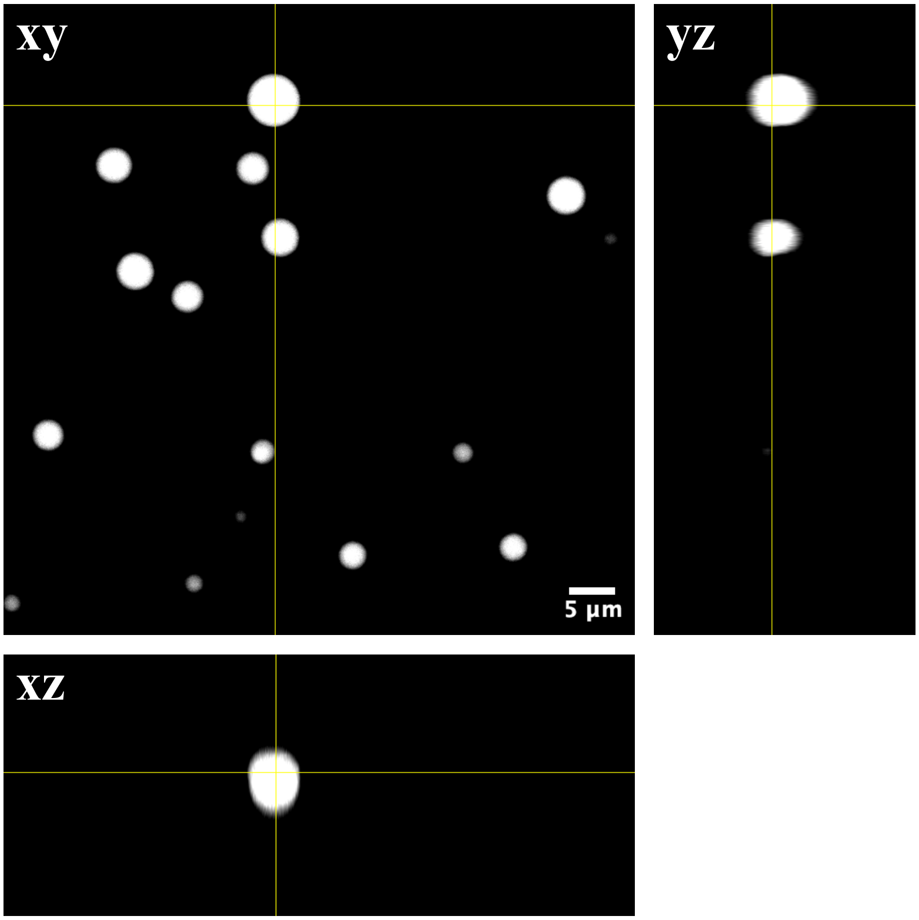


FIG. S2. Orthographic projections of representative z-stack image indicating droplets are mostly spherical in shape.

##### e. FRAP image analysis

The raw images contain many droplets. We first (i) identify a region of interest (ROI) containing only the bleached condensate and then (ii) compute the mean pixel intensity and radius for that condensate. (i) Using the image processing module from the *SciPy* library, we first applied a Gaussian filter ($\sigma = 1.15$) to denoise the images immediately before and after bleaching, and then subtracted the latter pixel intensities from the former over the entire image array to obtain the difference image. The difference image was segmented using Otsu's method, which establishes a pixel intensity threshold by maximizing inter-class variance [6]. We reduced this threshold by 20% to include more of the condensate periphery. Following hole filling and morphological closing for continuity, the condensate mask was applied to the difference image to estimate the condensate centroid and radius. The ROI containing the condensate was then defined as a square region with side length 4 × radius, centered at the identified condensate centroid. (ii) The ROI in the prebleach image was segmented using the same Otsu's method to obtain the final condensate mask, from which we computed the mean pixel intensity, centroid, area ($A$), and circular radius ($\sqrt{A/\pi}$). Using the same segmentation protocol on the recovery images, we tracked the droplet centroid in each frame and translated the condensate mask from the prebleach image to align with the droplet centroid in each frame. These mask shifts were necessary to account for droplet drift within the focal plane during imaging. The shifted masks were then used to extract raw mean fluorescence intensity curves for the measured droplets.

The raw intensities were normalized using:

$$I_{\mathrm{norm}}(t) = \frac{I_{\mathrm{raw}}(t) - I_{\mathrm{min}}}{I_{\mathrm{max}} - I_{\mathrm{min}}}, \tag{S45}$$

where $I_{\mathrm{raw}}(t)$ is the raw fluorescence intensity of a droplet at a given time $t$, $I_{\mathrm{min}}$ is the minimum intensity (i.e., the fluorescence immediately after bleaching), and $I_{\mathrm{max}}$ is the maximum intensity (i.e., the fluorescence immediately before bleaching). The normalized intensity $I_{\mathrm{norm}}(t)$ for each measured droplet was then fitted as described in the earlier section to extract the characteristic timescale $\tau$.

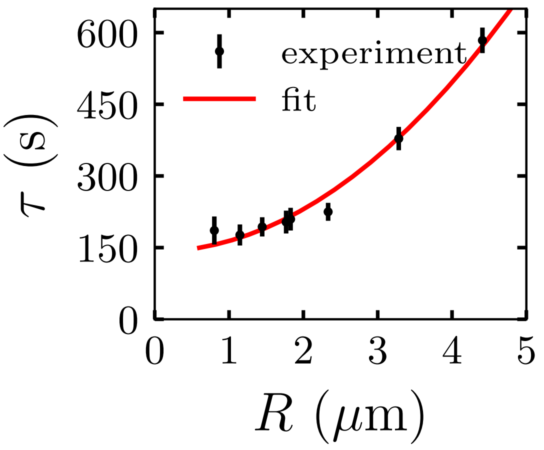


FIG. S3. Exchange timescale versus droplet radius for droplets measured in a replicated experiment, fitted with a shifted quadratic function of the form $\tau = a + bR^2$, where $a = 141.7 \pm 10.2$ s and $b = 22.1 \pm 1.2\,\mu\mathrm{m}^{-2}$s. Error bars on the data points represent $3\sigma$.

#### f. Reproducibility of shifted quadratic scaling

To verify the shifted quadratic trend reported in the main text, we conducted an independent set of whole-droplet FRAP experiments over the same size range ($\sim 1\,\mu$m to $5\,\mu$m) on a different sample batch prepared on a separate day. Plotting the timescale versus droplet size, we again observed that the same scaling law held (Fig. S3). This is consistent with the findings reported in the main text, which attribute the origin of the shifted quadratic scaling to a switch from dense-phase diffusion-limited recovery in large droplets ($R \gtrsim 2\,\mu$m) to bulk conversion-limited recovery in smaller droplets.

* Contact authors: (RK); (YZ)

[1] J. Crank, *The Mathematics of Diffusion*, Oxford science publications (Clarendon Press, 1979).
